# riboWaltz: optimization of ribosome P-site positioning in ribosome profiling data

**DOI:** 10.1101/169862

**Authors:** Fabio Lauria, Toma Tebaldi, Paola Bernabò, Ewout J.N. Groen, Thomas H. Gillingwater, Gabriella Viero

## Abstract

Ribosome profiling is a powerful technique used to study translation at the genome-wide level, generating unique information concerning ribosome positions along RNAs. Optimal localization of ribosomes requires the proper identification of the ribosome P-site in each ribosome protected fragment, a crucial step to determine trinucleotide periodicity of translating ribosomes, and draw correct conclusions concerning where ribosomes are located. To determine the P-site within ribosome footprints at nucleotide resolution, the precise estimation of its offset with respect to the protected fragment is necessary. Here we present riboWaltz, an R package for calculation of optimal P-site offsets, diagnostic analysis and visual inspection of data. Compared to existing tools, riboWaltz shows improved accuracies for P-site estimation and neat ribosome positioning in multiple case studies.

**Availability and Implementation:** riboWaltz was implemented in R and is available at https://github.com/LabTranslationalArchitectomics/RiboWaltz

## Introduction

Ribosome profiling (RiboSeq) is an experimental technique used to investigate translation at single nucleotide resolution and genome-wide scale (Ingolia et al., 2009; Ingolia et al., 2012), through the identification of short RNA fragments protected by ribosomes from nuclease digestion (Steitz et al., 1969; Wolin et al., 1988). The last few years have witnessed a rapid adoption of this technique and a consequent explosion in the volume of RiboSeq data (Michel and Baranov 2013; Brar and Weissman, 2015). In parallel, a number of dedicated computational algorithms were developed for extracting transcript-level information, including novel translation initiation sites, coding regions and differentially translated genes (Xiao et al., 2016; Zhong et al., 2017), as well as positional information describing fluxes of ribosomes along the RNA at sub-codon resolution (Martens et al., 2015, Legendre et al., 2016) and conformational changes in ribosomes during the elongation step of translation (Lareau et al., 2014).

Much of this information relies on the ability to determine, within ribosome protected fragments (reads), the exact localization of the P-site, i.e. the site holding the t-RNA, which is linked to the growing polypeptide chain during translation. This position can be specified by the distance of the P-site from both 5’ and 3’ ends of the reads, the so-called P-site Offset, PO (**Figure 1A**). Accurate determination of the PO is a crucial step to verify the trinucleotide periodicity of ribosomes along coding regions (Ingolia et al., 2009, Guo et al., 2010), derive reliable translation initiation and elongation rates (Gritsenko et al., 2015; Michel et al., 2014), accurately estimate codon usage bias and translation pauses (Pop et al., 2014, Weinberg et al., 2016), and reveal novel translated regions in known protein coding transcripts or ncRNAs (Hsu et al., 2016; Kochetov et al., 2016; Raj et al., 2016).

**Figure 1.**
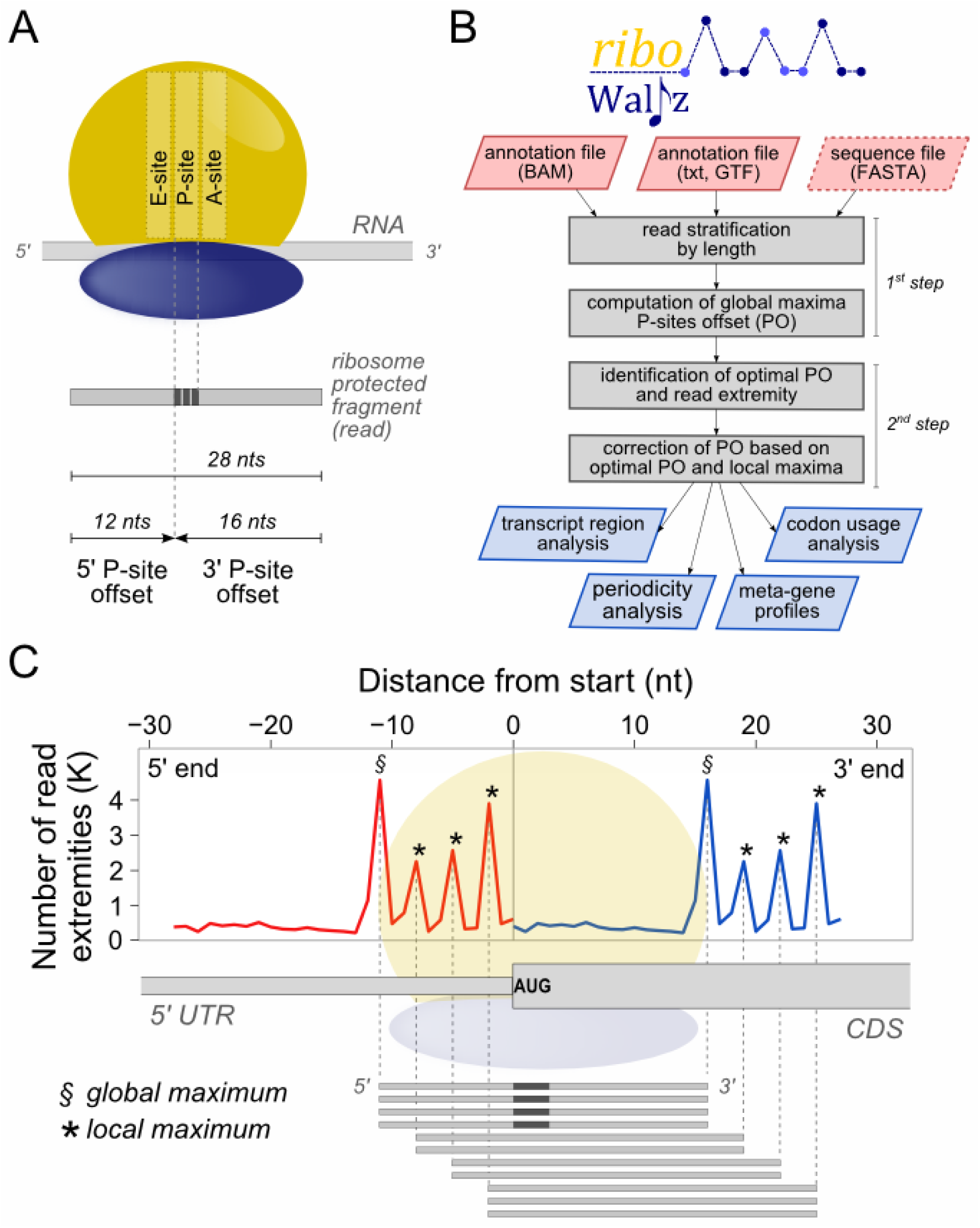
(**A**) Schematic representation of the P-site offset. Two offsets can be defined, one for each extremity of the read. (**B**) Flowchart representing the basic steps of riboWaltz, its inputs requirements and outputs. (**C**) An example of ribosome occupancy profile obtained from the alignment of the 5’ and the 3’ end of reads around the start codon (reads length, 28 nucleotides) is superimposed to the schematic representation of a transcript, a ribosome positioned on the TIS and a set of reads used for generating the profiles.

Typically the PO is defined as a constant number of nucleotides from either the 3′ or 5′ end of ribosome protected fragments, independently from their length (**Figure 1A**) (Gao et al., 2015). This approach may lead to an inaccurate detection of the P-site’s position owing to potential offset variations associated with the length of the reads. This problem is frequently resolved by selecting subsets of reads with defined length (Bazzini et al., 2014; Han et al., 2014). As such, this procedure removes from the analysis reads that are potentially derived from fragments associated to alternative conformations of the ribosome (Chen et al., 2012; Budkevich et al., 2014) and characterized by shorter or longer lengths (Lareau et al., 2014). Recently, computational tools have been developed to assist with RiboSeq analysis and P-site localization, for example Plastid (Dunn and Weissman, 2016) and RiboProfiling (Popa et al., 2016). Both tools compute the PO after stratifying the reads in bins, according to their length. However, each bin is treated independently, possibly leading to excessive variability of the offsets across bins.

Here, we describe the development of riboWaltz, an R package aimed at computing the PO for all reads from single or multiple RiboSeq samples. Taking advantage of a two-step algorithm where offset information is passed through populations of reads with different length in order to maximize offset coherence, riboWaltz computes with extraordinary precision the PO, showing higher accuracy and specificity of P-site positions than the other methods. riboWaltz provides the user with a variety of graphical representations, laying the foundations for further accurate RiboSeq analyses and better interpretation of positional information.

## Implementation

### Input acquisition and processing

riboWaltz requires two mandatory input data: a set of BAM files from transcriptome alignments of one or multiple RiboSeq samples and a text file with minimal transcript annotations, including the length of coding sequences and UTRs of known protein coding transcripts (**Figure 1B**). Optionally, a third file containing sequence information (in fasta format) can be provided as input to perform P-site specific codon sequence analysis. riboWaltz acquires BAM files and converts them into BED files utilizing the *bamtobed* function of the BEDTools suite (Quinlan and Hall, 2010).

### Identification of the P-site position

The identification of the P-site (defined by the position of its first nucleotide within the reads) is based on reads aligning across annotated translation initiation sites (TIS or start codon), and in particular on the distance between their extremities and the start codon itself, as proposed by Ingolia et al., 2009.

riboWaltz specifically infers the PO for each sample in two-steps. At first, riboWaltz groups by length (L) the reads mapping on TIS. To avoid biases in PO calculation, reads whose extremities are too close to the start codon, identified by a parameter called “flanking length” (FL), are discarded from further analysis. Then, for each length group, riboWaltz generates the occupancy profiles of read extremities, i.e. the number of 5’ and 3’ read ends in the region around the start codon (**Figure 1C**). For each length group, we defined temporary 5’ and 3’ POs (tPO) as the distance between the first nucleotide of the TIS and the nucleotide corresponding to the global maximum found in the profiles of the 5’ and the 3’ end at the left and at the right of the start codon, respectively (**Figure 1C**). Therefore, considering the occupancy profiles as a function *f* of the nucleotide position *x* with respect to the TIS, the temporary 5’ and 3’ PO for reads of length (L) are such that:

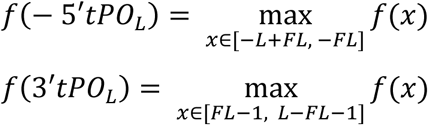

The two sets of length-specific temporary POs are defined as:

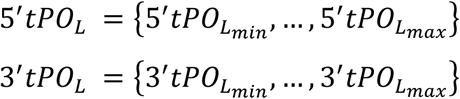

where *L_min_* and *L_max_* are respectively the minimum and the maximum length of the reads. At the end of this first step, the temporary POs are applied to all the reads (R), obtaining two sets of read-specific tPOs:

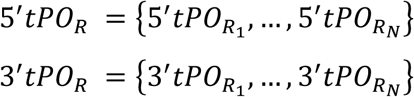

where N is the number of reads.

Despite good estimation of P-site positions, artifacts may arise from either the small number of reads with a specific length or the presence of noise in the signal, potentially producing inaccurate results. In other words, the offset estimated independently from the global maximum of each read length is not necessarily the best choice. This approach can produce high variability in PO values of reads differing for only one nucleotide in length (See **Supplementary Tables 1-3**) To minimize this problem, riboWaltz performs a second step for correcting the temporary POs.

The most frequent PO (called optimal PO, oPO) and the associated extremity (optimal extremity) are chosen as reference points to adjust the other values. The optimal PO is selected between the two modes of read specific tPO sets ( *Mode*(5′*tPO_R_*) and *Mode*(3′*tPO_R_*)) as the one with the highest frequency.

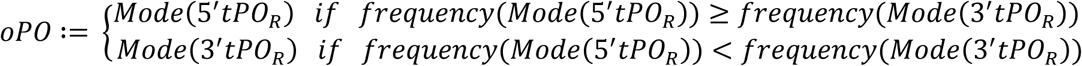

where the notation | | indicates the cardinality of a set.

Note that this step also selects the optimal extremity to calculate the corrected PO. The correction step is read-length-specific and works as follows: if the offset associated to a length bin is equal to the optimal PO no changes are made. Otherwise, i) the local maxima of the occupancy profiles are extracted; ii) the distances between the first nucleotide of the TIS and each local maxima is computed; iii) the new PO is defined as the distance in point ii) that is closest to the optimal PO. Summarizing, given the local maxima position (LMP) of the occupancy profile for the optimal extremity, the corrected PO for reads of length L (*cPO_L_*) is such that

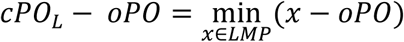

Finally, the corrected POs are applied to all the reads.

### Output

riboWaltz returns three data structures. The first is a list of sample-specific data frames containing for each read,, i) the position of the P-site (identified by the first nucleotide of the codon) with respect to the beginning of the transcript; ii) the distance between the P-site and both the start and the stop codon of the coding sequence; iii) the region of the transcript (5′ UTR, CDS, 3′ UTR) where the P-site is located iv) optionally, if a sequence file is provided as input, the sequence of the triplet covered by the P-site. The second data structure is a data frame reporting the percentage of reads aligning across the start codon (if any) and on the whole transcriptome, stratified by sample and read length. Moreover, this file includes the P-site offsets before and after the optimization (tPO and cPO values). The third data structure is a data frame containing, for each transcript, the number of ribosome protected fragments with in-frame P-site mapping on the CDS. This data frame can be used to estimate transcript-specific translation levels and perform differential analysis comparing multiple conditions.

riboWaltz also provides several graphical outputs, described in more detail in the Results section. All graphical outputs are returned as lists containing an object of class ggplot (further customizable by the user), and a data frame containing the source data for the plots.

## Results

### riboWaltz overview

In order to show riboWaltz functionalities, we analyzed a ribosome profiling dataset obtained from mouse brains (authors’ unpublished data, see **Supplementary Methods**).

riboWaltz integrates several graphical functions that provide multiple types of output results. First, the distribution of the length of the reads (**Figure 2A**): this is a useful preliminary inspection tool to understand the contribution of each length to the final P-site determination, and possibly decide to remove certain lengths from further analyses. Second, the percentage of P-sites located in the 5’ UTR, CDS and 3’ UTR regions of mRNAs compared with a uniform distribution weighted on region lengths, simulating random P-site positioning along mRNAs (**Figure 2B**). This analysis is a good way to verify the expected enrichment of ribosome signal in the CDS. Third, to understand if, and to which extent, P-site determination results in codon periodicity in the CDS, riboWaltz produces a plot withthe percentage of P-sites matching one of the three possible translation reading frames (phase analysis) for 5’ UTR, CDS and 3’ UTR, stratifying reads by length (**Figure 2C**). Fourth, the meta-gene read density heatmap, based on the position of read extremities and stratifying reads by length (**Figure 2D**). This plot provides an overview of the occupancy profiles used for P-site determination and allows to check by visual inspection if PO values are reasonable and possibly proceed with manual modification. Fifth, to understand which codons, if any, present higher or lower density of ribosome protected fragments, riboWaltz provide the user with the analysis of the empirical codon usage, i.e. the frequency of in-frame P-sites along the coding sequence associated to each codon, normalized for codon frequency in sequences (**Figure 2E**). Indeed, the comparison of these values in different biological conditions can be of great help to unravel possible defects in aa-tRNAs use or ribosome elongation at specific codons. Finally, single transcripts profiles and meta-gene profiles based on P-site position can be generated (**Figure 3B, top row**) with multiple options: i) combining multiple replicates applying convenient scale factors provided by the user, ii) considering each replicate separately, or iii) stratifying the reads by length.

**Figure 2.**
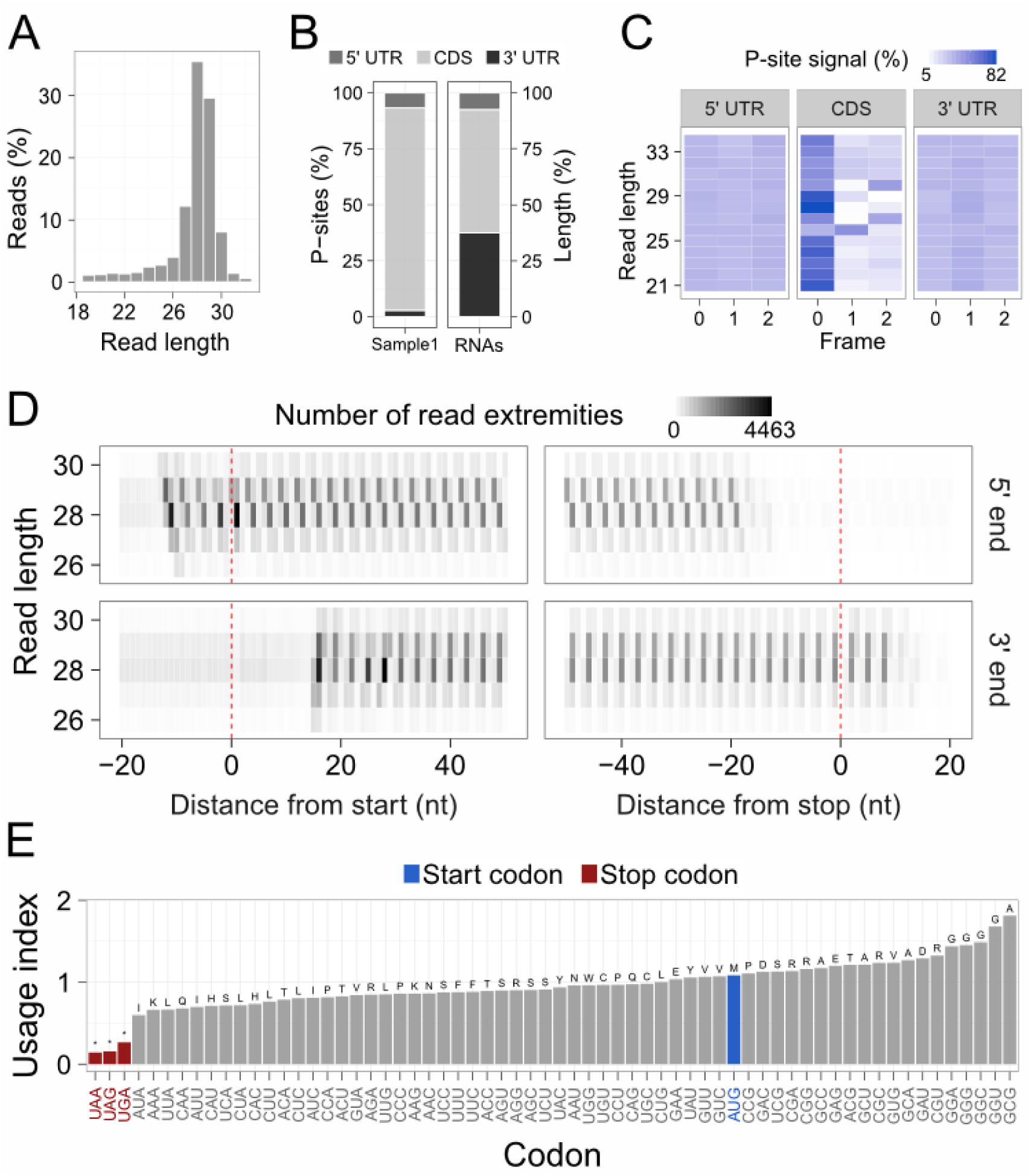
(**A**) Distribution of the read lengths. (**B**) Left, percentage of P-sites in the 5’ UTR, CDS and 3’ UTR of mRNAs from ribosome profiling data. Right, percentage of region lengths in mRNAs sequences. (**C**) Percentage of P-sites in the three frames along the 5’ UTR, CDS and 3’ UTR, stratified for read length. (**D**) Example of meta-gene heatmap reporting the signal associated to the 5’ end (upper panel) and 3’ end (lower panel) of the reads aligning around the start and the stop codon for different read lengths. (**E**) Codon usage analysis based on in-frame P-sites. Codon usage index is calculated as the frequency of in-frame P-sites along the coding sequence associated to each codon, normalized for codon frequency in sequences. Aminoacids corresponding to each codon are displayed above each bar. All panels were obtained from ribosome profiling of whole mouse brain.

### Comparison with other tools

We tested riboWaltz on three ribosome profiling datasets in yeast (*S. cerevisiae*, Lareau et al., 2014), mouse brain (authors’ unpublished data) and human cell lines (Hek-293, Gao et al., 2015) and compared our results to those obtained using RiboProfiling (v1.2.2, Popa et al., 2016) and Plastid (v0.4.5, Dunn and Weissman, 2016) (**Figure 3** and **Supplementary Tables 1-3**). For RiboWaltz and RiboProfiling, the percentage of P-sites with correct frame within the CDS region was comparable in all cases, while Plastid showed lower performances (**Figure 3A** and **Supplementary Figure 1A and 2A**). Remarkably, meta-profiles produced by riboWaltz displayed a neat periodicity uniquely in the CDS (**Figure 3B** and **Supplementary Figure 1B** and **2B**), with almost no signal along UTRs, neither in the proximity of the start nor of the stop codon. By contrast, Plastid and RiboProfiling generated a shift of the start of the periodic region toward the 5’ UTR (**Figure 3B** and **Supplementary Figure 1B** and **2B**), suggesting a possible mislocalization of ribosomes before the start and stop codons (**Supplementary Figure 1B and 2B**), an issue that has the potential to generate inaccurate biological conclusions.

**Figure 3.**
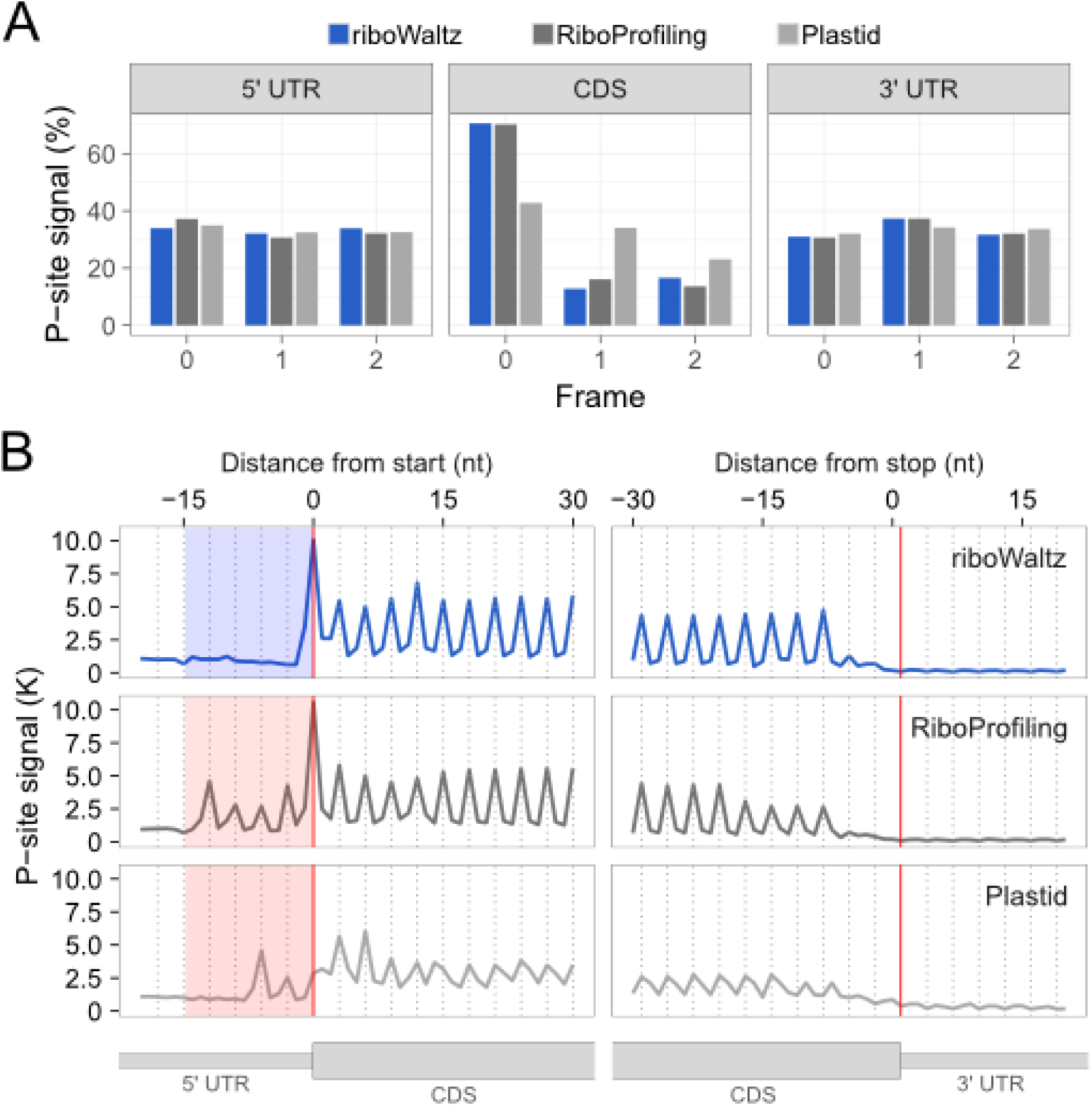
(**A**) Percentage of P-sites in the three frames along the 5’ UTR, CDS and 3’ UTR from ribosome profiling performed in mouse brain and (**B**) meta-profiles showing the periodicity of ribosomes along transcripts at genome-wide scale, based on P-site identification by riboWaltz, RiboProfiling and Plastid. The shaded areas to the left of the start codon highlight the shift of the periodicity toward the 5’ UTR that is absent in the case of data analysed using riboWaltz.

## Conclusions

In conclusion, riboWaltz identifies with high precision the position of ribosome P-sites from ribosome profiling data. By improving on other currently-available approaches, riboWaltz can assist with the detailed interrogation of RiboSeq data at single nucleotide resolution, providing precise information that may lay the groundwork for further positional analyses and new biological discoveries.

## Acknowledgements

We thank the Core Facility, Next Generation Sequencing Facility (HTS) CIBIO, University of Trento (Italy) for technical support.

## Supplementary Information

**Supplementary Figure 1.**
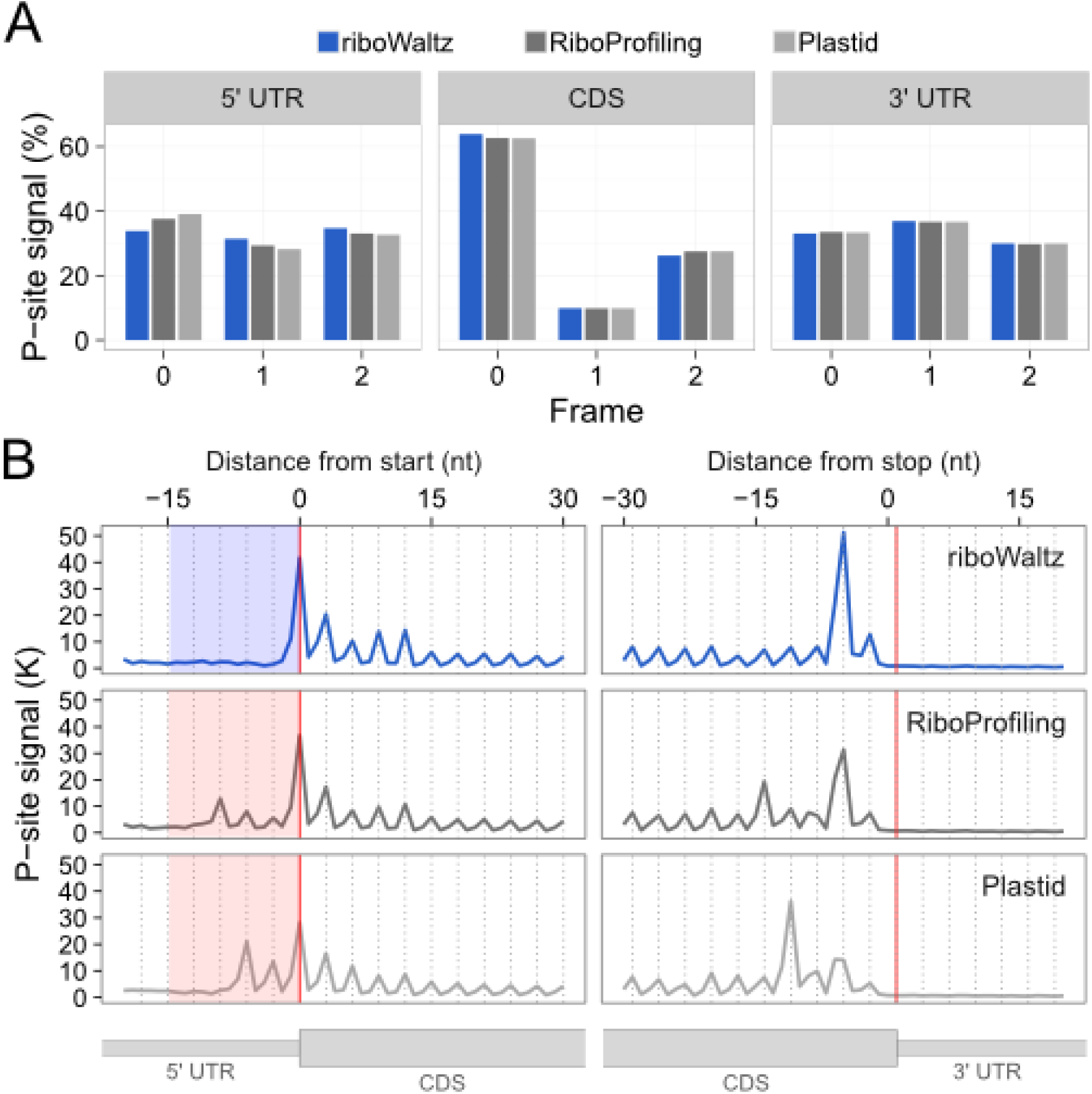
(**A**) Percentage of P-sites in the three frames along the 5’ UTR, CDS and 3’ UTR from ribosome profiling in Hek-293 (Gao et al., 2015) and (**B**) meta-profiles showing the periodicity of ribosomes along transcripts at genome-wide scale, based on P-site identification by riboWaltz, RiboProfiling and Plastid. The shaded areas to the left of the start codon highlight the shift of the periodicity toward the 5’ UTR.

**Supplementary Figure 2.**
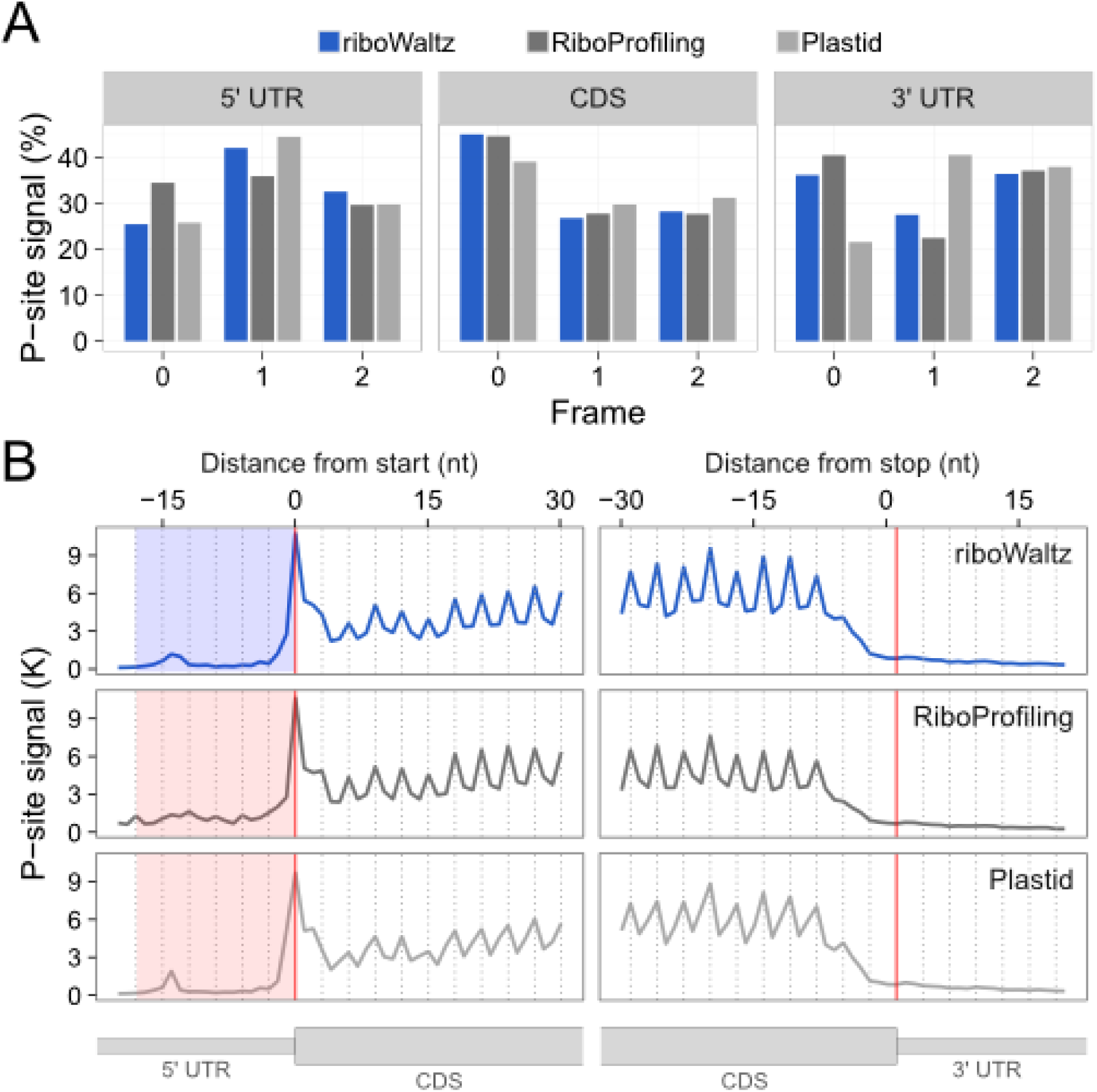
(**A**) Percentage of P-sites in the three frames along the 5’ UTR, CDS and 3’ UTR from ribosome profiling in yeast (*S.* cerevisiae, Lareau et al., 2014) and (**B**) meta-profiles showing the periodicity of ribosomes along transcripts at genome-wide scale, based on P-site identification by riboWaltz, RiboProfiling and Plastid. The shaded areas to the left of the start codon highlight the shift of the periodicity toward the 5’ UTR.

**Supplementary Table 1:**
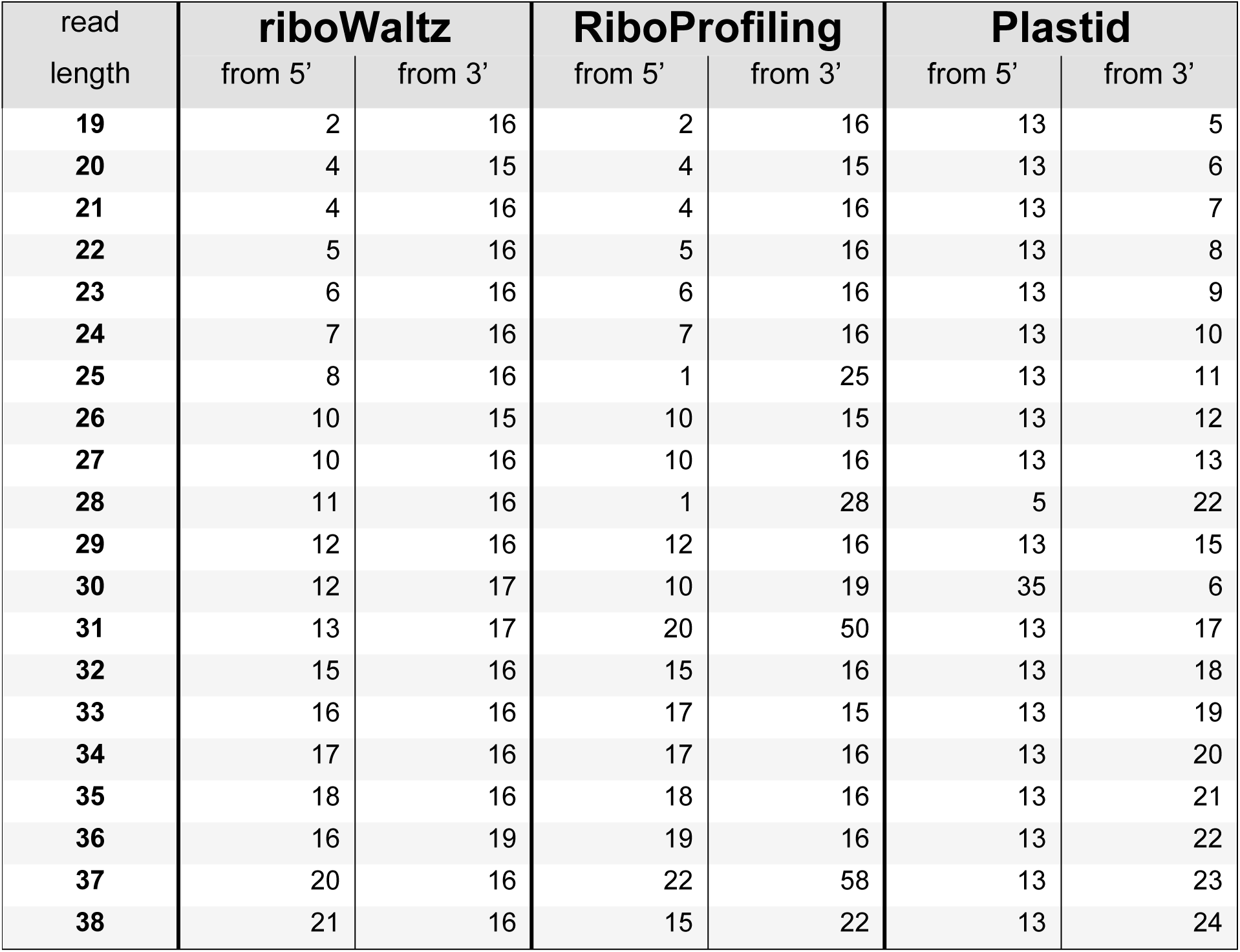
Comparison of the P-site offsets identified for each read length by riboWaltz, RiboProfiling and Plastid in mouse. The PO computed from both read extremities are reported.

**Supplementary Table 2:** Comparison of the P-site offsets identified for each read length by riboWaltz, RiboProfiling and Plastid in human. The PO computed from both read extremities are reported.

| read<br>length | **riboWaltz** |  | **RiboProfiling** |  | **Plastid** |  |
| --- | --- | --- | --- | --- | --- | --- |
|  | from 5' | from 3' | from 5' | from 3' | from 5' | from 3' |
| **25** | 12 | 12 | 0 | 24 | 3 | 21 |
| **26** | 12 | 13 | 12 | 13 | 12 | 13 |
| **27** | 12 | 14 | 12 | 14 | 12 | 14 |
| **28** | 12 | 15 | 3 | 24 | 6 | 21 |
| **29** | 12 | 16 | 12 | 16 | 6 | 22 |
| **30** | 12 | 17 | 12 | 17 | 12 | 17 |
| **31** | 13 | 17 | 9 | 21 | 9 | 21 |
| **32** | 13 | 18 | 10 | 21 | 7 | 24 |
| **33** | 12 | 20 | 12 | 20 | 50 | 18 |
| **34** | 15 | 18 | 17 | 16 | 13 | 20 |

**Supplementary Table 3:** Comparison of the P-site offsets identified for each read length by riboWaltz, RiboProfiling and Plastid in yeast. The PO computed from both read extremities are reported.

| read<br>length | riboWaltz |  | RiboProfiling |  | Plastid |  |
| --- | --- | --- | --- | --- | --- | --- |
|  | from 5' | from 3' | from 5' | from 3' | from 5' | from 3' |
| 21 | 12 | 8 | 12 | 8 | 12 | 8 |
| 22 | 13 | 8 | 50 | 71 | 13 | 8 |
| 23 | 13 | 9 | 2 | 20 | 13 | 9 |
| 24 | 13 | 10 | 22 | 45 | 13 | 10 |
| 25 | 13 | 11 | 9 | 15 | 13 | 11 |
| 26 | 12 | 13 | 44 | 69 | 13 | 12 |
| 27 | 13 | 13 | 10 | 36 | 13 | 13 |
| 28 | 12 | 15 | 12 | 15 | 12 | 15 |
| 29 | 13 | 15 | 13 | 15 | 12 | 16 |
| 30 | 12 | 17 | 12 | 17 | 12 | 17 |
| 31 | 13 | 17 | 13 | 17 | 13 | 17 |
| 32 | 14 | 17 | 14 | 17 | 13 | 18 |
| 33 | 14 | 18 | 43 | 75 | 13 | 19 |
| 34 | 15 | 18 | 3 | 36 | 13 | 20 |
| 35 | 10 | 24 | 5 | 39 | 13 | 21 |
| 36 | 13 | 22 | 11 | 24 | 13 | 22 |
| 37 | 15 | 21 | 12 | 48 | 13 | 23 |
| 38 | 14 | 23 | 23 | 60 | 13 | 24 |
| 39 | 22 | 16 | 12 | 26 | 13 | 25 |
| 40 | 7 | 32 | 7 | 32 | 13 | 26 |

## Supplementary methods

### RiboSeq data processing

Raw reads were processed by removing 5’ adapters, discarding reads shorter than 20 nucleotides and trimming the first nucleotide (using Trimmomatic v0.36). Reads mapping on rRNAs and tRNAs (downloaded from the SILVA rRNA and the Genomic tRNA databases respectively) were removed. The remaining reads were aligned to the organism transcriptome with Bowtie2 (v2.2.6) employing the default settings. All reads aligning to the very same region were collapsed to avoid potential PCR duplicates, and only strand-specific reads were kept.

## References

Bazzini, A. A., Johnstone, T. G. Christiano, R., Mackowiak, S. D. Obermayer, B., Fleming, E. S. … & Giraldez, A. J. (2014). Identification of small ORFs in vertebrates using ribosome footprinting and evolutionary conservation. The EMBO journal, e201488411.

Brar, G. A., & Weissman, J. S. (2015). Ribosome profiling reveals the what, when, where and how of protein synthesis. Nature Reviews Molecular Cell Biology.

Budkevich, T. V., Giesebrecht, J., Behrmann, E., Loerke, J., Ramrath, D. J., Mielke, T., … & Sanbonmatsu, K. Y. (2014). Regulation of the mammalian elongation cycle by subunit rolling: a eukaryotic-specific ribosome rearrangement. Cell, 158(1), 121–131.

Chen, J., Tsai, A., O’Leary, S. E., Petrov, A., & Puglisi, J. D. (2012). Unraveling the dynamics of ribosome translocation. Current opinion in structural biology, 22(6), 804–814.

Dunn, J. G., & Weissman, J. S. (2016). Plastid: nucleotide-resolution analysis of next-generation sequencing and genomics data. BMC genomics, 17(1), 958.

Gao, X., Wan, J., Liu, B., Ma, M., Shen, B., & Qian, S. B. (2015). Quantitative profiling of initiating ribosomes in vivo. Nature methods, 12(2), 147–153.

Gritsenko, A. A., Hulsman, M., Reinders, M. J. & de Ridder, D. (2015). Unbiased Quantitative Models of Protein Translation Derived from Ribosome Profiling Data. PLoS Comput Biol, 11(8), e1004336.

Guo, H., Ingolia, N. T., Weissman, J. S., & Bartel, D. P. (2010). Mammalian microRNAs predominantly act to decrease target mRNA levels. Nature, 466(7308), 835–840.

Han, Y., Gao, X., Liu, B., Wan, J., Zhang, X., & Qian, S. B. (2014). Ribosome profiling reveals sequence-independent post-initiation pausing as a signature of translation. Cell research, 24(7), 842–851.

Hsu, P. Y., Calviello, L., Wu, H. Y. L., Li, F. W., Rothfels, C. J. Ohler, U., & Benfey, P. N. (2016). Super-resolution ribosome profiling reveals unannotated translation events in Arabidopsis. Proceedings of the National Academy of Sciences, 113(45), E7126–E7135.

Ingolia, N. T., Brar, G. A., Rouskin, S., McGeachy, A. M. & Weissman, J. S. (2012). The ribosome profiling strategy for monitoring translation in vivo by deep sequencing of ribosome-protected mRNA fragments. Nature protocols, 7(8), 1534–1550.

Ingolia, N. T., Ghaemmaghami, S., Newman, J. R., & Weissman, J. S. (2009). Genome-wide analysis in vivo of translation with nucleotide resolution using ribosome profiling. Science, 324(5924), 218–223.

Kochetov, A. V., Allmer, J., Klimenko, A. I., Zuraev, B. S., Matushkin, Y. G., & Lashin, S. A. (2016). AltORFev facilitates the prediction of alternative open reading frames in eukaryotic mRNAs. Bioinformatics, btw736.

Legendre, R., Baudin-Baillieu, A., Hatin, I., & Namy, O. (2015). RiboTools: a Galaxy toolbox for qualitative ribosome profiling analysis. Bioinformatics, 31(15), 2586–2588.

Lareau, L. F., Hite, D. H., Hogan, G. J., & Brown, P. O. (2014). Distinct stages of the translation elongation cycle revealed by sequencing ribosome-protected mRNA fragments. Elife, 3, e01257.

Martens, A. T. Taylor, J., & Hilser, V. J. (2015). Ribosome A and P sites revealed by length analysis of ribosome profiling data. Nucleic acids research, gkv200

Michel, A. M., & Baranov, P. V. (2013). Ribosome profiling: a Hi-Def monitor for protein synthesis at the genome-wide scale. Wiley Interdisciplinary Reviews: RNA, 4(5), 473–490.

Michel, A. M., Andreev, D. E., & Baranov, P. V. (2014). Computational approach for calculating the probability of eukaryotic translation initiation from ribo-seq data that takes into account leaky scanning. BMC bioinformatics, 15(1), 380.

Pop, C., Rouskin, S., Ingolia, N. T., Han, L., Phizicky, E. M., Weissman, J. S., & Koller, D. (2014). Causal signals between codon bias, mRNA structure, and the efficiency of translation and elongation. Molecular systems biology, 10(12), 770.

Popa, A., Lebrigand, K., Paquet, A., Nottet, N., Robbe-Sermesant, K., Waldmann, R., & Barbry, P. (2016). RiboProfiling: a Bioconductor package for standard Ribo-seq pipeline processing. F1000Research, 5.

Quinlan, A. R., & Hall, I. M. (2010). BEDTools: a flexible suite of utilities for comparing genomic features. Bioinformatics, 26(6), 841–842.

Raj, A., Wang, S. H., Shim, H., Harpak, A., Li, Y. I., Engelmann, B., … & Pritchard, J. K. (2016). Thousands of novel translated open reading frames in humans inferred by ribosome footprint profiling. Elife, 5, e13328.

Steitz, J. A. (1969). Polypeptide chain initiation: nucleotide sequences of the three ribosomal binding sites in bacteriophage R17 RNA. Nature, 224, 957–964.

Weinberg, D. E., Shah, P., Eichhorn, S. W., Hussmann, J. A., Plotkin, J. B., & Bartel, D. P. (2016). Improved ribosome-footprint and mRNA measurements provide insights into dynamics and regulation of yeast translation. Cell reports, 14(7), 1787–1799.

Wolin, S. L., & Walter, P. (1988). Ribosome pausing and stacking during translation of a eukaryotic mRNA. The EMBO journal, 7(11), 3559.

Xiao, Z., Zou, Q., Liu, Y., & Yang, X. (2016). Genome-wide assessment of differential translations with ribosome profiling data. Nature communications, 7.

Zhong, Y., Karaletsos, T., Drewe, P., Sreedharan, V. T., Kuo, D., Singh, K., … & Rätsch, G. (2017). RiboDiff: detecting changes of mRNA translation efficiency from ribosome footprints. Bioinformatics, 33(1), 139–141.

